# White Tea Mitigates Obesity-Related Metabolic Disorders Through Apelin and Insulin Modulation in Rats

**DOI:** 10.1101/2024.10.05.616820

**Authors:** Ayşegül Sümer, Öznur Demirtaş, Esra Pınarbaş, Eda Yılmaz Kutlu, Hülya Kılıç

## Abstract

Obesity, a prevalent global health issue, is influenced by genetic, metabolic, environmental, and behavioral factors, often leading to serious complications like type 2 diabetes, cardiovascular diseases, and reduced life expectancy. Recent studies have highlighted apelin, a bioactive peptide, for its significant role in obesity and related metabolic disorders. Apelin, primarily found in adipose tissue, regulates appetite, energy expenditure, and insulin sensitivity, making it a promising target for antiobesity treatments. Concurrently, tea, especially white tea, rich in bioactive compounds, has shown potential health benefits, including antiobesity effects. This study investigates the impact of white tea on weight gain, glucose metabolism, and lipid metabolism by analyzing its effects on apelin-13, apelin-36, and insulin levels in a high-fat diet (HFD)-induced obesity model in rats. Thirty-two male Sprague Dawley rats were divided into four groups: control, HFD, HFD+orlistat, and HFD+white tea. Over 13 weeks, the HFD groups received their respective treatments while being monitored for weight gain and metabolic changes. Results indicated that the HFD group exhibited significant weight gain and elevated cholesterol levels compared to the control. White tea-treated rats showed reduced weight gain and lower cholesterol levels, suggesting its antiobesity properties. Apelin-13 levels were significantly lower in the HFD group, while white tea mitigated this effect, highlighting its potential role in modulating apelin levels and improving insulin sensitivity. The findings support the hypothesis that white tea can be an effective antiobesity agent, offering a safer, natural alternative to conventional treatments. Further research is warranted to elucidate the precise mechanisms underlying these beneficial effects.

## Introduction

Obesity is a chronic condition marked by the accumulation and enlargement of fat cells within adipose tissue, primarily resulting from an imbalance where caloric intake surpasses energy expenditure. It has become a prevalent global health issue, influenced by a complex interplay of genetic, metabolic, environmental, and lifestyle factors. This condition elevates the risk of numerous health complications, including type 2 diabetes, cardiovascular diseases, cancer, and disorders affecting the musculoskeletal system. Obesity can also lead to impaired life quality and reduced life expectancy (3, 4). Therefore, protective and/or preventive alternative lifestyle changes have become interesting.

Apelin, a naturally occurring peptide that acts as the endogenous ligand for the APJ receptor, has garnered significant interest due to its multifaceted physiological roles, especially in regulating cardiovascular functions, fluid balance, and energy metabolism. Recent studies underscore its pivotal role in the pathogenesis of obesity and related metabolic disorders. Apelin is expressed in a number of different tissues, including adipose tissue, where it modulates a number of crucial processes, including appetite regulation, energy expenditure and insulin sensitivity. It has been demonstrated that apelin levels are frequently elevated in individuals with obesity and correlate with body mass index (BMI) and insulin resistance. Furthermore, apelin’s central and peripheral actions make it a promising target for therapeutic interventions aimed at ameliorating obesity and its associated complications. Thus, a deeper understanding of the intricate role of apelin in obesity may facilitate the development of novel strategies for combating this global health challenge. (5-8)

An increasing body of evidence suggests that tea, particularly green and white varieties, which are rich in bioactive compounds, possesses significant preventive and therapeutic potential against various diseases. These teas are particularly noted for their robust protective effects on health (9).

Tea, derived from the Camellia sinensis plant, has been integral to human culture for millennia and remains a popular beverage across the globe, including in Turkey. This natural herb is composed of various primary components, including polyphenols, amino acids, alkaloids, sugars, proteins, pectin, aromatic compounds, enzymes, and organic acids. Previous research has reported that tea contains beneficial active components with various medical benefits, including anticancer, antioxidant, anti-inflammatory, antibacterial, cardiovascular protection, anti-sugar, and antiobesity properties (10). Studies have shown the positive effects of tea consumption, which is especially rich in polyphenols such as epigallocatechin gallate (EGCG), on energy balance, body weight and glucose metabolism (11). However, high doses of EGCG, which is highly abundant in tea, have been reported to show cytotoxic and hepatotoxic effects in rodents (11,12). These results raise the question of whether crude extracts from tea or brewed tea could be a more effective and safer, easy-to-consume alternative antiobesity strategy. Green, white, yellow, oolong, black, and dark teas are the most common types of tea

White tea is produced from freshly harvested shoots and young leaves, with minimal processing. This results in a tea with high antioxidant levels, although it is oxidised by at least 5% (13). White tea, which has many medicinal effects due to its minimal processing, has been reported to be associated with CVD, diabetes, cancer, metabolism due to the strong antioxidant activities of phenolic substances in its content (14)

*Camellia sinensis* is the primary agricultural crop cultivated in Rize, Turkey. The biological effects of other tea types (green, black, oolong) and their ingredients have been the subject of extensive research. Similar studies have been conducted on white tea, yet the mechanism underlying its antiobesity effect remains unclear. The number of studies examining the effects of apelin and white tea on obesity is limited. This study aimed to investigate the impact of white tea on weight gain, glucose metabolism and lipid metabolism by examining its effects on apelin-13, apelin 36 and insulin levels in a high-fat diet-induced obesity model. Additionally, this study sought to elucidate one of the mechanisms underlying the antiobesity effect of white tea.

## Materials and Methods

### Chemicals

White tea was provided from the General Directorate of Tea Enterprises (ÇAYKUR). Orlistat, 120 mg capsule Xenical, Roche S.p.A., Segrate (MI), Italy), rat apelin-13 Enzyme-linked Immunosorbent Assay (ELISA) kit (Bt Lab, Cat No. E1427Ra, Lot:202103014) rat apelin-36 ELISA kit (Bt Lab, Cat No. E1331Ra, Lot:202103014), rat insülin ELISA kit (Bt Lab, Cat No. E0707Ra, Lot:202103014),, High fat diet (Arden Research & Experiment, 22% kcal from fat)(Table 1), Ketamine (Ketalar, 100mg/10mL, Pfizer) and Xylazine Hydrochloride (Rompun, 2% 25mL, Bayer).

**Table 1.**
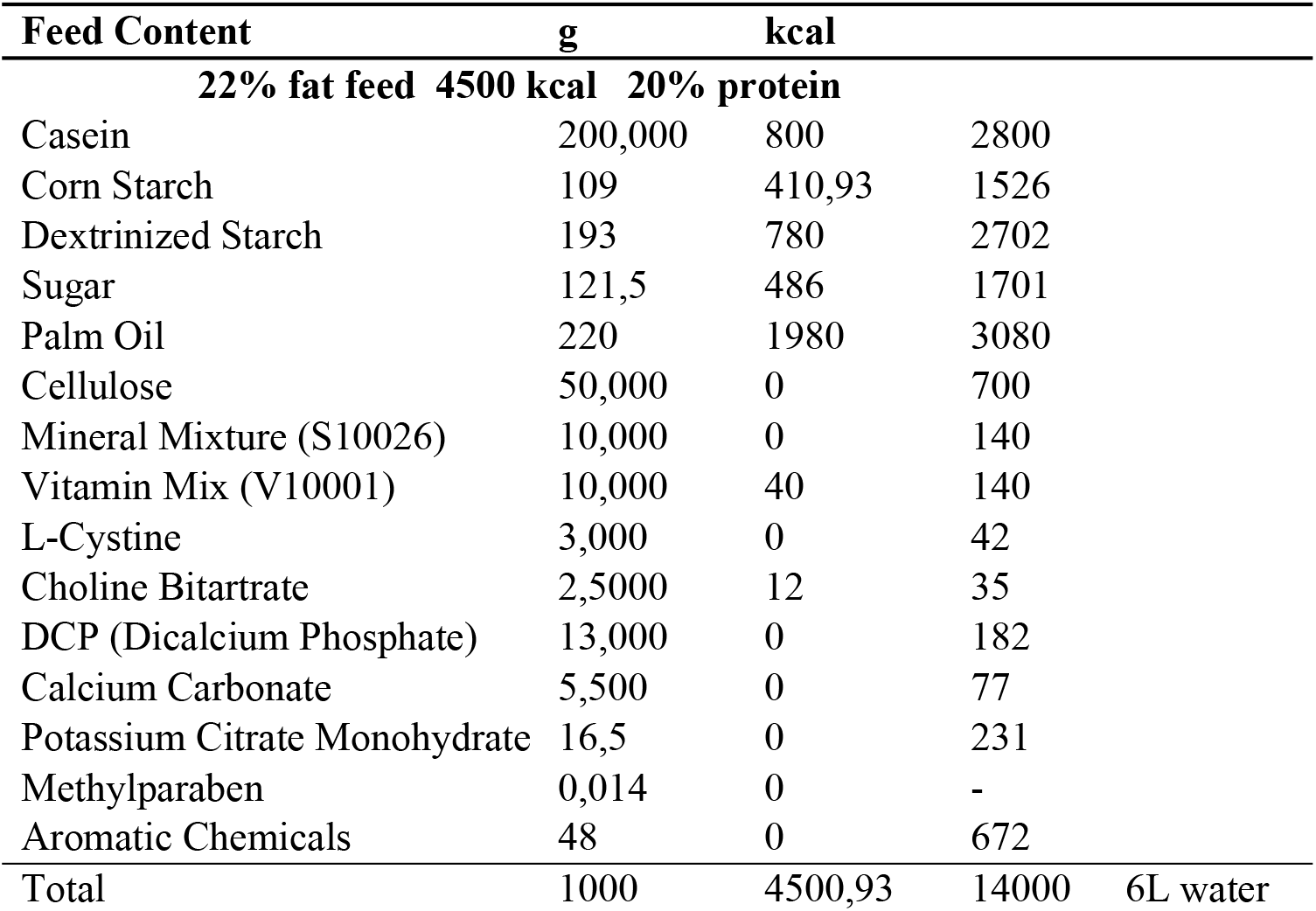
High fat dietary content.

### Animals and procedures

This study received approval from the Local Ethics Committee for Animal Experiments at Recep Tayyip Erdoğan University (RTEU) under protocol number 2020/39. The experiment involved 32 male Sprague Dawley rats, aged 6 to 8 weeks, sourced from the RTEU Experimental Animal Research and Application Centre. The rats were maintained at a temperature of 23°C ± 2°C, humidity of 55% ± 5%, and a 12-hour day/night cycle. The cages were cleaned on a weekly basis, and sawdust was utilized as litter. The rats were provided with a standard diet (Bayramoğlu Yem ve Un San. Tic.A.Ş., Full Pellet Rat Feed) ad libitum for one week and acclimated to the laboratory environment. They were then randomly selected and divided into four groups of eight rats (n=8) per group, housed in transparent polyethylene cages. In order to create an obesity model, a high-fat diet (HFD, 22% kcal from fat) was provided ad libitum at a rate of 15-20 g/day per rat. The animals were provided with water from specially designed bottles. At the conclusion of the experiment, one rat from the control group and one from the orlistat group were lost. The experiments, statistical analyses, and calculations were based on these data points.

### Tea and Orlistat Preparation

The harvesting of white tea occurs on an annual basis, commencing in the spring. All studies were conducted using the same harvest. Fresh white tea is prepared on a daily basis using the appropriate brewing technique. The dose was prepared at a concentration of 5 mg per kg for each rat, with 1 mL administered orally to each rat daily. As this is an experimental model of obesity, it was anticipated that the weight of the rats would increase during the course of the experiment. The rats were weighed on a weekly basis, and the white tea solutions were prepared in accordance with the desired concentration.

To illustrate, for a 250 g rat, a 1 mL gavage application was prepared by weighing 1.25 mg of white tea, adding it to 1 mL of boiled water, and allowing it to stand for 10 minutes. This solution was then administered to the rat. Given the minute volume and weight of the solutions, they were prepared by measuring out 10 mL (15). The rats in the HFD group were subjected to dietary and other treatments until their weight exceeded that of the rats in the C group by more than 20% (16).

Orlistat (Xenical) is a weight-loss medication that inhibits pancreatic lipase, thereby blocking the digestion of triglycerides and reducing the absorption of dietary fat (17). In the study, Orlistat was dissolved in distilled water at a dose of 30 mg/kg and administered daily to the rats in the group via oral gavage.The dose calculation was updated and applied according to the changing weights of the rats. For example, for a 250 g rat, 7.5 mg orlistat was weighed and dissolved in 1 mL distilled water and administered to the rats by gavage.

### Biochemical Experiments

The animals were fasted overnight, and blood samples were obtained from the intracardiac left ventricle under anaesthesia with ketamine and xylazine hydrochloride. The animals were administered hydrochloride at a ratio of 3:1. The levels of apelin-13, apelin-36, and insulin (I) were determined using the ELISA method. Triglyceride (TG), total cholesterol (TC), high-density lipoprotein cholesterol (HDL-C), and low-density lipoprotein cholesterol (LDL-C) concentrations and fasting blood glucose (FBG) levelswere quantified using the Beckman Coulter AU5800 autoanalyzer. The Homeostatic Model Assessment for Insulin Resistance (HOMA-IR) was calculated using the following formula and was used for the detection of insulin resistance.

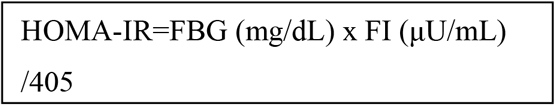

### Statistics

The data analysis and evaluation were conducted using the SPSS 25 package program. To assess whether the numerical data followed a normal distribution, skewness and kurtosis were calculated. Data were considered normally distributed if the skewness and kurtosis values were within the range of −3 to +3 The Kruskal-Wallis test and the post hoc adjusted Bonferroni test were employed to ascertain the existence of a difference between the groups. The results were expressed as a median and the 25th to 75th percentile (interquartile range, IQR). Spearman’s correlation analysis was employed to assess the strength of the relationship between the variables. The threshold for statistical significance was set at p < 0.05.

## Results

### Weight measurements

At the conclusion of the 13-week study period, the mean weight gain rates for the C, HFD, HFD+OL, and HFD+WT groups were 53%, 82%, 102.5%, and 68%, respectively. The successful establishment of the obesity model is evidenced by the fact that the final weight of rats in the HFD group was 20% higher than in the C group. A statistically significant difference was observed between the final body weights of the HFD group and the C and HFD+WT groups (X2 = 18.340, p = 0.000) (Table 2).

**Table 2.** Final body weights of the groups.

| | n | Median | IQR | $\chi^2$ | p |
| --- | --- | --- | --- | --- | --- |
| **Body weight (g)** |  |  |  |  |  |
| Control (1) | 7 | 460.00 | 50.00 |  |  |
| HFD (2) | 8 | 549.50 | 23.50 |  |  |
| HFD+OL (3) | 7 | 478.00 | 81.00 | 18.340 | 0.000* |
| HFD+ WD (4) | 8 | 423.50 | 44.75 |  |  |

*IQR: Interquartile Range, $\chi^2$ : Kruskal-Wallis Test, HFD: High Fat Diet, HFD+OL: High Fat Diet +Orlistat, HFD+WT: High Fat Diet +White Tea, $p<0.05$ , \*Bonferroni 4<2, 1<2*

## Biochemical Results

### Insulin, Apelin 13, Apelin 36 Levels

A significant reduction in apelin 13 levels was observed in the high-fat diet (HFD) group in comparison to the control group. (X^2^=9.965, p=0.019), while no statistically significant difference was observed between the apelin-36 levels of the groups (p>0.05) (Table 3). A Kruskal-Wallis test was employed to ascertain whether there were any statistically significant differences in the median values of insulin, apelin 13, and apelin 36 levels between the groups. A statistically significant difference was observed between the medians of apelin 13 levels in the groups (X2 = 9.965, p = 0.019). The adjusted Bonferroni method was employed to ascertain which group exhibited the discrepancy. It was determined that the apelin 13 levels of the high-fat diet group were lower than those of the control group. No statistically significant difference was identified between the insulin and apelin 36 levels of the groups (p > 0.05).

**Table 3.**
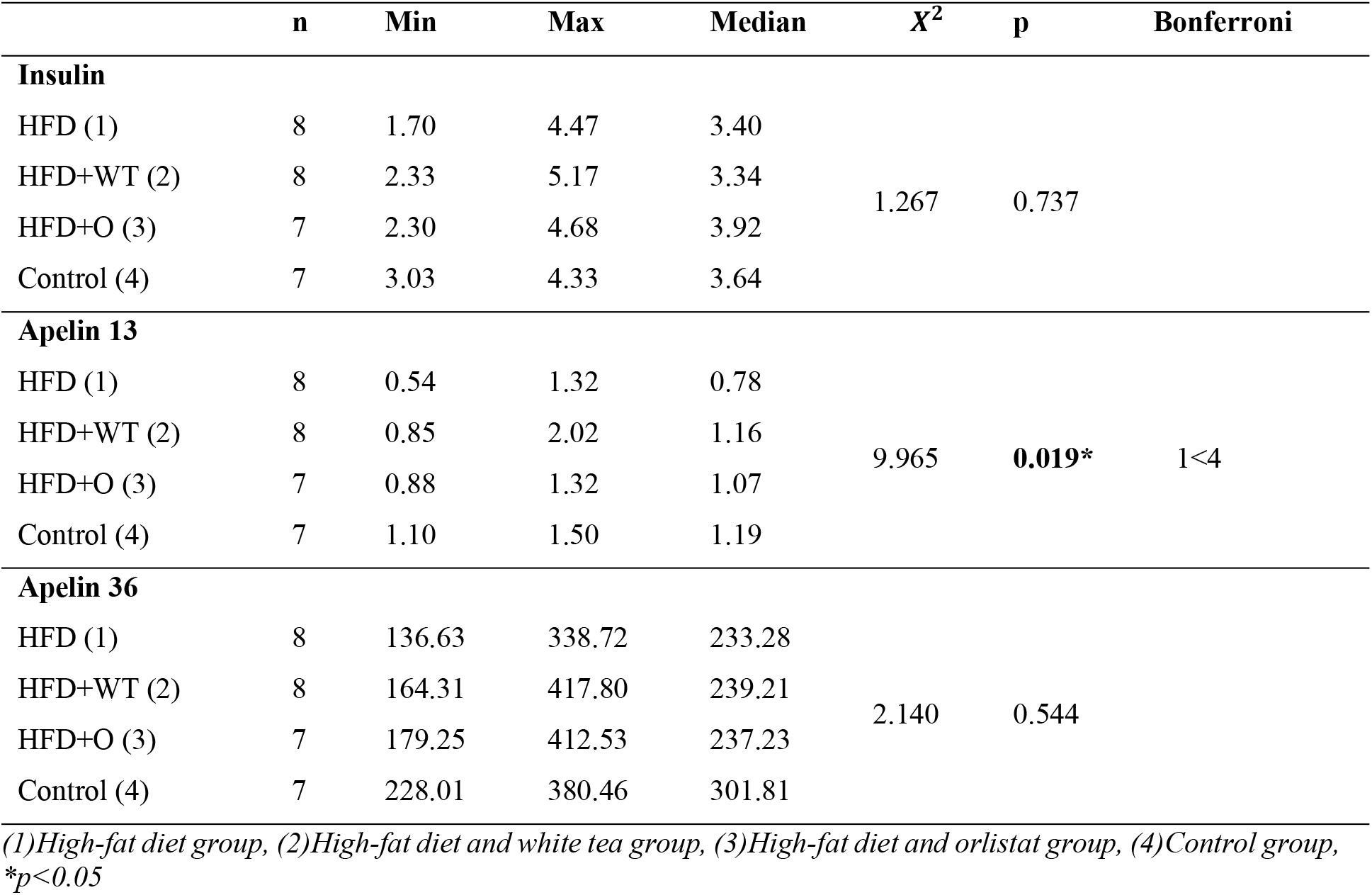
Insulin, apelin 13, apelin 36 levels of the groups.

|  | n | Min | Max | Median | X<sup>2</sup> | p | Bonferroni |
| --- | --- | --- | --- | --- | --- | --- | --- |
| Insulin |  |  |  |  |  |  |  |
| HFD (1) | 8 | 1.70 | 4.47 | 3.40 | 1.267 | 0.737 |  |
| HFD+WT (2) | 8 | 2.33 | 5.17 | 3.34 |  |  |  |
| HFD+O (3) | 7 | 2.30 | 4.68 | 3.92 |  |  |  |
| Control (4) | 7 | 3.03 | 4.33 | 3.64 |  |  |  |
| Apelin 13 |  |  |  |  |  |  |  |
| HFD (1) | 8 | 0.54 | 1.32 | 0.78 | 9.965 | 0.019* | 1<4 |
| HFD+WT (2) | 8 | 0.85 | 2.02 | 1.16 |  |  |  |
| HFD+O (3) | 7 | 0.88 | 1.32 | 1.07 |  |  |  |
| Control (4) | 7 | 1.10 | 1.50 | 1.19 |  |  |  |
| Apelin 36 |  |  |  |  |  |  |  |
| HFD (1) | 8 | 136.63 | 338.72 | 233.28 | 2.140 | 0.544 |  |
| HFD+WT (2) | 8 | 164.31 | 417.80 | 239.21 |  |  |  |
| HFD+O (3) | 7 | 179.25 | 412.53 | 237.23 |  |  |  |
| Control (4) | 7 | 228.01 | 380.46 | 301.81 |  |  |  |

(1)High-fat diet group, (2)High-fat diet and white tea group, (3)High-fat diet and orlistat group, (4)Control group, \* $p < 0.05$

### HOMA-IR Results

There was no statistically significant difference observed among the groups regarding insulin and HOMA-IR levels (p > 0.05). However, glucose levels showed a statistically significant difference between the C group and the HFD group (X^2^=8.824, p=0.32) (Figure 1). Additionally, analysis of the relationships between apelin-13, apelin-36, insulin, and HOMA-IR levels revealed a statistically significant positive correlation between insulin and HOMA-IR (r = 0.455, p < 0.05). On the other hand, no significant correlation was found between the other parameters (p > 0.05) (Table 4).

**Table 4.** Correlation between apelin-13, apelin-36, insulin and HOMA-IR levels.

| Parameters |  | Apelin-13 | Apelin-36 | Insulin | HOMA-IR |
| --- | --- | --- | --- | --- | --- |
| **Apelin-13** | r | 1 | 0.237 | 0.179 | -0.344 |
|  | p | - | 0.207 | 0.345 | 0.063 |
| **Apelin-36** | r |  | 1 | 0.301 | 0.033 |
|  | p |  | - | 0.106 | 0.863 |
| **Insulin** | r |  |  | 1 | 0.455 |
|  | p |  |  | - | 0.012* |
| **HOMA-IR** | r |  |  |  | 1 |
|  | p |  |  |  | - |

*r=Spearman's korelasyon, \*p<0.05*

**Figure 1.**
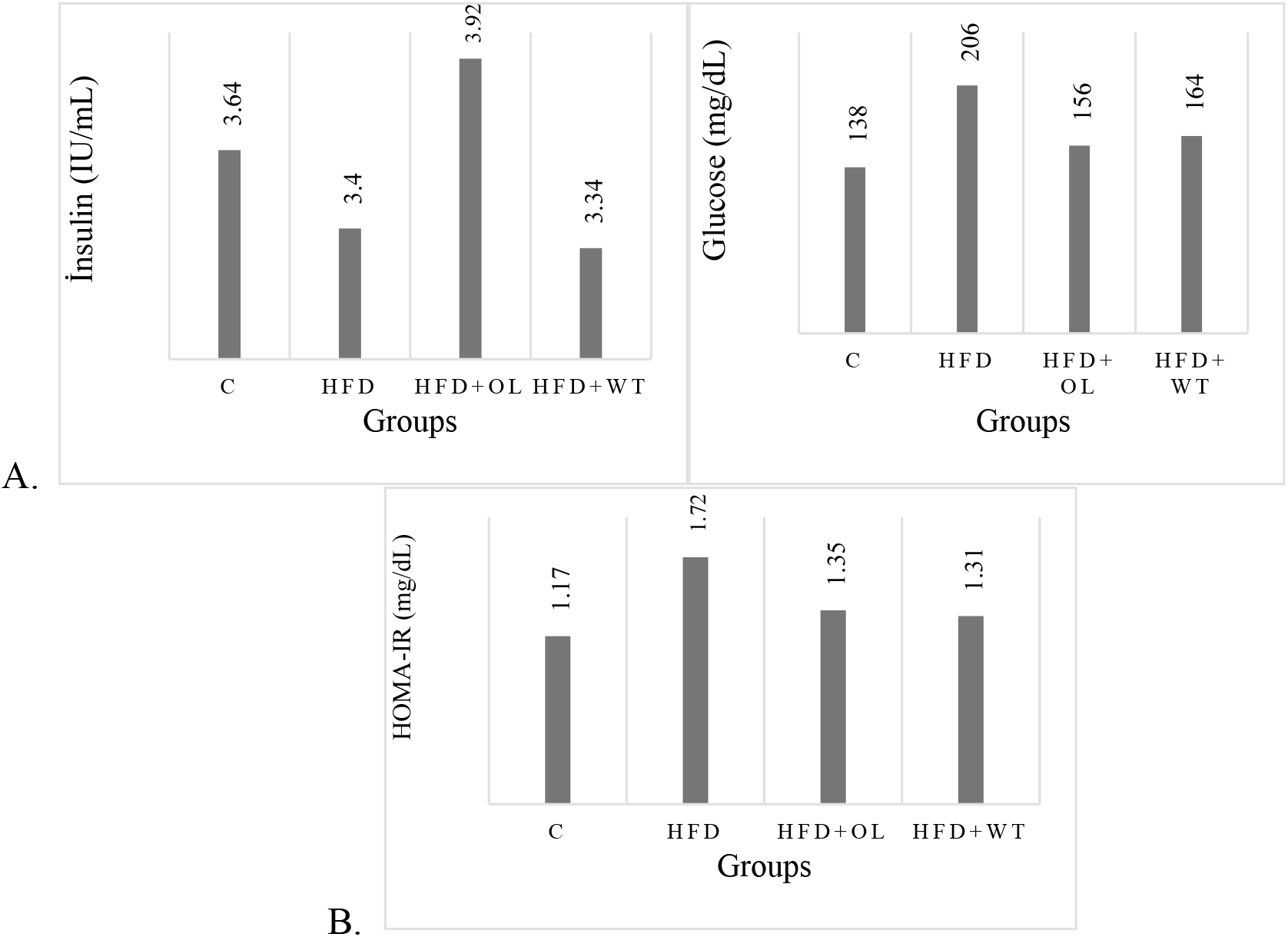
A. Insulin,and Glucose and B. HOMA-IR levels of the groups

### Lipid Parameters

The control group exhibited significantly lower total cholesterol levels compared to the HFD group (X^2^=9.435, p=0.024). Additionally, there were no statistically significant differences found among the groups regarding TG, HDL-C, and LDL-C levels (p > 0.05) (Table 5).

**Table 5.** Lipid parameters of the groups.

| | n | Median | IQR | $\chi^2$ | p |
| --- | --- | --- | --- | --- | --- |
| Cholesterol (mg/dL) |  |  |  |  |  |
| HFD (1) | 8 | 64.00 | 15.25 | 9.435 | 0.024* |
| HFD+WT (2) | 8 | 53.50 | 14.25 |  |  |
| HFD+OL (3) | 7 | 57.00 | 15.00 |  |  |
| Control (4) | 7 | 49.00 | 9.00 |  |  |
| Triglyceride (mg/dL) |  |  |  |  |  |
| HFD (1) | 8 | 27.00 | 22.75 | 1.191 | 0.755 |
| HFD+WT (2) | 8 | 28.50 | 10.75 |  |  |
| HFD+OL (3) | 7 | 34.00 | 17.00 |  |  |
| Control (4) | 7 | 35.00 | 12.00 |  |  |
| HDL-C (mg/dL) |  |  |  |  |  |
| HFD (1) | 8 | 40.85 | 8.50 | 6.623 | 0.085 |
| HFD+WT (2) | 8 | 34.85 | 8.80 |  |  |
| HFD+OL (3) | 7 | 33.70 | 10.70 |  |  |
| Control (4) | 7 | 33.30 | 2.90 |  |  |
| LDL-C (mg/dL) |  |  |  |  |  |
| HFD (2) | 8 | 16.50 | 12.50 | 7.615 | 0.055 |
| HFD+WT (2) | 8 | 14.50 | 5.50 |  |  |
| HFD+OL (3) | 7 | 16.00 | 6.00 |  |  |
| Control (4) | 7 | 9.00 | 8.00 |  |  |

*Min: Minimum, Max: Maximum, IQR; Interquartile Range, $\chi^2$ : Kruskal-Wallis Test, (1)High-fat diet group, (2)High-fat diet and white tea group, (3)High-fat diet and orlistat group, (4)Control group $p<0.05$ , \*Bonferroni $1<2$*

## Discussion

This study aimed to investigate the effects of white tea on obesity by measuring apelin-13 and apelin-36, insulin, glucose, HOMA-IR, TC, TG, HDL-C, and LDL-C in an obesity model induced by a high-fat diet. Although the Sprague Dawley breed is not optimal for use in an obesity model, a 20% increase in body weight was observed in the high-fat diet (HFD) groups compared to the control group, which is considered a criterion for obesity.

It is established that white tea possesses antiobesity properties (18, 19). The results of this study indicated that the orlistat group exhibited the greatest weight gain, followed by the high-fat diet group, the white tea group, and the control group, in that order. The final body weight and adipose tissue increase of the rats was the highest in the HFD group, followed by the HFD+OL, C, and HFD+WT groups, respectively. The higher weight gain observed in the orlistat group may be attributed to the administration of the drug in the morning. Given that rats are nocturnal active animals and that orlistat should be taken just before a meal, it was concluded that the results were acceptable. Although the orlistat group exhibited the greatest weight gain, the highest final body weight was observed in the HFD group, indicating that adipose tissue expansion was more pronounced in this group. It was hypothesised that the lowest weight gain and final body weight in the control group were attributable to the highest initial body weight of the control group. The growth of adipose tissue was the least in the HFD+WT group, and the final body weight was significantly lower than that of the HFD group. This indicates that white tea has an antiobesitic effect, which is in line with the findings of previous studies. Given that the white tea application was conducted in the morning, it can be posited that the observed effect is indicative of a stimulation of the metabolic rate, a reduction in the formation of adipose tissue and an induction of lipolysis.

The antidiabetic effects of tea are well documented and have been demonstrated in numerous studies (19, 20). Similarly, the data revealed a statistically significant difference in glucose levels between the groups, with the highest levels observed in the HFD group, as anticipated. (p < 0.05). No significant difference was observed between the insulin values of the groups. (p>0.05). The highest HOMA-IR levels were observed in the HFD group, indicating that white tea may possess hypoglycaemic properties by reducing blood sugar and enhancing glucose tolerance. Additionally, white tea may diminish insulin resistance and/or augment insulin sensitivity, rather than increasing insulin secretion.

The precise mechanisms through which white tea exerts its therapeutic effects remain unclear. However, evidence suggests that it may be effective in the treatment of obesity and its associated complications by alleviating lipid metabolism disorders (21). A reduction in lipid metabolism parameters was observed in rats treated with white tea (22). A study investigating the hypolipidemic effects of white tea demonstrated that white tea has potent lipolytic and anti-adipogenic effects on human subcutaneous preadipocytes (23).

The study revealed that the highest cholesterol values were observed in the HFD group, followed by the HFD+OL, HFD+WT and control groups. A statistically significant difference was identified between the groups (p<0.05). These findings lend support to the hypothesis that white tea has hypocholesterolemic effects, as LDL-C levels were found to be highest in the high-fat diet group and lowest in the white tea group. No statistically significant difference was observed between the triglyceride (TG) and high-density lipoprotein cholesterol (HDL-C) levels of the groups (p > 0.05). It was hypothesised that the elevated lipid parameters observed in the orlistat group were attributable to the administration of orlistat in the morning.

The findings of apelin levels in physiological and pathological situations in studies have been influenced by a number of factors, including ethnicity, gender, age groups, body fat distribution, different isoforms of apelin and the various methods applied. (24)

In a study comparing HFD-induced obese mice with the control group, it was shown that apelin and insulin levels were not significantly different between the groups, but there was a strong correlation between apelin and insulin in the obese and control mouse groups. It has been concluded that apelin and insulin interact independently of obesity. (25). Apelin is upregulated in obesity. In clinical and experimental studies, it has been observed that serum apelin level or adipose tissue expression increases in cases of obesity and insulin resistance (26, 27). A study in obese young individuals reported that Apelin levels have been decreased, especially in prediabetic patients, compared to the control group (27). In the early stages of metabolic disorders such as obesity, prediabetes or newly diagnosed type 2 DM, apelin expression decreases due to hyperinsulinemia. As the duration of hyperglycemia increases, apelin expression increases in various metabolic disorders such as deterioration of insulin resistance and insulin secretion due to damage to β cells. (28).

The Apelin 13 levels were found to be the lowest in the HFD group, followed by the orlistat, white tea and control groups, respectively (p < 0.05). Furthermore, apelin-36 levels exhibited a parallel trend with apelin 13 levels, with no statistically significant difference (p > 0.05). The observed decrease in Apelin levels in the HFD group may be attributed to the increase in adipose tissue associated with obesity and the complications thereof, including hyperglycaemia, hypercholesterolemia and hyperinsulinemia. It can be proposed that elevated apelin levels in obesity and associated disorders represent a compensatory mechanism for reducing the effects of obesity. The absence of an expected increase in apelin levels in the HFD group suggests that the macroscopically observed increase in adipose tissue did not affect the systemic level and did not cause insulin resistance. Despite the successful creation of obesity in rats through the control of weight gain, it was concluded that the negative metabolic and systemic effects of obesity did not fully manifest. It was hypothesised that the reason for this might be the type of animal used and the composition of the feed. The beneficial effects of white tea on lipid and carbohydrate metabolism indicate that it may result in less change in apelin levels in the white tea-treated group.While it has been found that the physiological effects of white tea may indirectly cause less change in apelin levels, it has been shown that white tea has antiobesity, hypoglycemic, insulin tolerance enhancing and hypocholesterolemia effects, and obesity and/or metabolic disorders affect apelin levels.

